# A highly rifampicin resistant *Mycobacterium tuberculosis* strain emerging in Southern Brazil

**DOI:** 10.1101/2020.09.03.282194

**Authors:** Maria Lucia Rossetti, Pedro Eduardo Almeida da Silva, Richard Steiner Salvato, Ana Júlia Reis, Sun Hee Schiefelbein, Andrea von Groll, Regina Bones Barcellos, Raquel Maschmann, Leonardo Esteves, Fernanda Spies, Rubia Raubach Trespach, Elis Regina Dalla Costa, Hermes Luís Neubauer de Amorim

## Abstract

Here we described phenotypical, molecular and epidemiological features of a highly rifampicin-resistant *Mycobacterium tuberculosis* strain emerging in Southern Brazil, that carries an uncommon insertion of 12 nucleotides at the codon 435 in the *rpoB* gene. Employing a whole-genome sequencing-based study on drug-resistant *Mycobacterium tuberculosis* strains, we identified this emergent strain in 16 (9.19%) from 174 rifampicin-resistant clinical strains included. Nine of these 16 strains were available to minimum inhibitory concentration determination and for all of them was found a high rifampicin-resistance level (≥ to 32 mg/L). This high resistance level could be explained by structural changes into the RIF binding site of RNA polymerase caused by the insertions, and consequent low-affinity interaction with rifampicin complex confirmed through protein modeling and molecular docking simulations. Epidemiological investigation showed that most of the individuals (56.25%) infected by the studied strains were prison inmate individuals or that spent some time in prison. The phylogenomic approach revealed that all strains carrying on the 12 nucleotide insertion belonged to the same genomic cluster, evidencing a communal transmission chain involving inmate individuals and community. We stress the importance of tuberculosis genomic surveillance and the introduction of measures to interrupt the *Mycobacterium tuberculosis* transmission chain in this region.

## 1. INTRODUCTION

Tuberculosis (TB) remains a major public health emergency worldwide and current efforts to control the disease have been threatened by the high rates of drug-resistant TB (DR-TB) cases[1]. The overall success rate for TB treatment is 82%, but multidrug resistance (MDR-TB) - resistance to isoniazid (INH) and rifampicin (RIF), the two main anti-TB drugs - is associated with worse treatment outcome, dropping the cure rates to 60%[1]. RIF is one of the most important drugs used in the anti-TB treatment, due to its high bactericidal effects. RIF mechanism action consists of binding at RNA polymerase (RNAp) beta-subunit, encoded by *rpoB* gene, resulting in the inhibition of the bacterial mRNA transcription[2,3]. The occurrence of RIF-resistance in *Mycobacterium tuberculosis (M. tuberculosis*) is generally associated to single nucleotide substitutions at *rpoB* gene, mainly at rifampicin resistance determining region (RRDR), an 81-base pair region comprising from codon 426 to 452[4].

Previous studies conducted in Rio Grande do Sul, a high TB burden setting and largest state in Southern Brazil, described an uncommon insertion of 12 nucleotides (12nt) at the codon 435 in the *rpoB* gene among MDR *M. tuberculosis* strains. The 12nt insertion results in a duplication of four amino acids (QNNP) and was firstly described by Perizzolo et al., (2012)[5]. Following studies analyzing MDR strains from Rio Grande do Sul[6–9] also reported strains carrying on the 12nt insertion, and to our knowledge, this insertion was not described in any other region worldwide.

Hence, for an enhanced TB control management in Southern Brazil, is need the elucidation concerning molecular and phenotypic consequences of that 12nt insertion, as well as, comprehension on this strain spreading into population. For this propose, we screened RIF-resistant *M. tuberculosis* strains from Rio Grande do Sul State using a WGS-based population study approach, in order to identify strains carrying on the 12nt insertion. Afterward, we examine the phylogenetic relationship among these strains, the phenotypic consequences of the insertion into resistance level and biological cost, and also the *in silico* prediction to RIF binding effect.

## 2. METHODS

### 2.1. Sample collection and drug susceptibility testing (DST)

In total 16 *M. tuberculosis* strains carrying on the 12nt insertion at *rpoB* gene were identified on a WGS-based state-wide study that is being currently conducted in Rio Grande do Sul, Southern Brazil. The study included 174 RIF-resistant strains collected from 2011 to 2014, and 169 of them presented multidrug resistance. The *M. tuberculosis* clinical strains were from the State Central Laboratory (LACEN - Rio Grande do Sul), the reference laboratory in charge of performing drug susceptibility testing (DST) from TB cases notified statewide. In the studied period around 246 RIF-resistant cases were notified in Rio Grande do Sul according to the SITE-TB website (national database for DR-TB cases reported in Brazil). Thus, our analysis covered 70.73% (174/246) from RIF-resistant cases registered in the period.

The DST was performed at LACEN using the liquid BACTEC™ MGIT™ 960 SIRE Kit for the BACTEC Mycobacteria Growth Indicator Tube 960 (MGIT 960) system (Becton Dickinson Diagnostic Systems, Sparks, MD). The test was conducted for the following first-line anti-TB drugs: RIF (1.0 mg/L), INH (0.1 mg/L), ethambutol (EMB) (5.0 mg/L) and streptomycin (STR) (1.0 mg/L).

### 2.2. DNA extraction

*M. tuberculosis* genomic DNA was extracted from sputum culture in Lowenstein–Jensen solid medium using Cetyltrimethylammonium Bromide (CTAB) method, as described by Van Embden et al. (1993)[10].

### 2.3. Whole Genome Sequencing

The 174 RIF-resistant clinical strains belonging to our state-wide study were subjected to WGS. Paired-end sequencing (2 × 150 bp) was performed on an Illumina NextSeq machine using either a 300 cycle v2 mid output or high output kit (Illumina, Code FC-404-2003 or Code FC-404-2004) under standard Illumina^®^ procedure as previously described[9].

### 2.4. Bioinformatics analysis

Raw reads were submitted to a routine pipeline to genomic variants identification. First, sequence reads were examined using FastQC (v0.11.7) (http://www.bioinformatics.babraham.ac.uk/projects/fastqc) and trimmed using Trimmomatic (v0.33) (parameters: LEADING:3 TRAILING:3 SLIDINGWINDOW:4:20 MINLEN:36)[11]. Reads were mapped against *M. tuberculosis* H37Rv reference genome (GenBank accession number: NC_000962.3) under *BWA-MEM* (v0.7.16) algorithm. The quality of the resulting BAM file was checked using *Qualimap*[12] and *sambamba* (v0.6.8) was used to mark read duplicates[13]. *Samtools* (v1.9)[14] and GATK (v3.8)[15] were used to variant calling from sorted mapped sequences. Variants were filtered based on the following criteria: mapping quality ≥50, base alignment quality ≥23 and minimum read depth of 10. Variant functional annotation was performed with *snpEff* (v4.3)[16].

SNP-based *M. tuberculosis* lineages were determined using *TB-Profiler*[17,18] pipeline in command-line version (2.7.4). Phylogenetic analysis was performed using *snippy* pipeline v4.3.6 (https://github.com/tseemann/snippy) for variant calling and alignment of all core-genomes. Variants positions within PE/PPE genes or other repetitive regions associated with low mappability scores were removed. A maximum-likelihood phylogenetic tree was generated using the software RAxML (v8.10.12)[19], applying the generalized time-reversible (GTR) model and 1000 bootstrap replicates. The resulting tree was rooted using *M. canettii* (Genbank accession number: NC_019950.1). A minimum spanning tree was also generated using Phyloviz (http://online2.phyloviz.net/) being implemented the goeBURST algorithm[20]. *In silico* spoligotyping profiles were obtained using *SpoTyping* (v2.0)[21] and the spoligotyping patterns were assigned to lineage using the SITVIT2 web database (http://www.pasteur-guadeloupe.fr:8081/SITVIT2).

### 2.5. Minimum inhibitory concentration (MIC)

Due to the retrospective nature of this study, some strains were no longer available for additional phenotypic testing. Among the 16 strains carrying on the 12nt insertion, nine of them were viable to perform susceptibility tests. The minimum inhibitory concentration (MIC) to RIF for these nine available strains was determined using the resazurin microtiter assay (REMA) as previously described[22,23].

### 2.6. Determination of growth curve by Resazurin Reduction Method

The growth curve for two clinical strains with the 12nt insertion, three wild type clinical strains and one susceptible control strain (H37Rv) was determined by resazurin reduction method as previously reported by von Groll et al. (2010)[24]. Briefly, cultures were started in 96-well plate containing the bacterial inoculum in triplicate wells as well as the 7H9 broth alone as the blank control. After 48h incubated at 37°C, resazurin (viability indicator) was added to all test and blank control wells and re-incubated. The resazurin reduction (the blue indicator color turn to pink) was assessed by measuring every 12 hours the optical density (OD) at 620 nm with a plate reader (TECAN Spectrum Classic). Growth curves were constructed plotting the difference in OD between test and control wells every 12 hours.

The biological cost of each strain was estimated by the growth index (GI) which was the time needed by each strain to reach an OD of 0.4 starting at an OD of 0.2. This time was calculated from the growth curve considering that all strains were in the logarithmic phase of growth between the two OD values. The biological cost was determined by the fitness relative (FR) which is the ratio of the GI of the insertion in each *rpoB* strains in relation to the H37Rv and in relation to the wild-type. In this case, a biological cost was considered if the FR was > 1, since the 12 nt insertion has a slow growth in relation to the susceptible ones.

### 2.7. Protein insertion modeling and effects on drug binding

The wild-type RNAp (RNAp_wt) structural model was obtained from the crystallographic structure deposited in the RCSB Protein Data Bank[25,26] (PDB) (www.rcsb.org) under accession code 5ZX3 (resolution 2.75 Å)[27]. The 3D structural models of the two mutants forms of the RNAp were obtained by homology modelling using the SWISS-MODEL Workspace[28]. Only the structures of the beta subunits of each mutant (*rpoB* INS1 and *rpoB* INS2) were modeled. The beta subunit of the structure under accession code 5ZX3 of PDB was used as template. The final structures for each RNAp mutant, here denominated RNAp_INS1 (carrying *rpoB* INS1) and RNAp_INS2 (carrying *rpoB* INS2), were obtained by fitting the coordinates of the modeled subunits with the coordinates of the beta subunit of the 5ZX3 crystallographic structure. The coordinates of the beta subunit of the template structure were subsequently removed, keeping the rest of the polymerase unchanged. The stereochemical quality of the models was checked by the QMEANDisCo[29] and QMEAN[30] scoring functions, implemented on the SWISS-MODEL platform[31], and with the analysis tools implemented on the MolProbity platform[32].

Docking simulations were carried out with two programs, AudoDock Vina (Vina)[33], version 1.1.2, and AutoDock[34], version 4.2. These programs use different algorithms and approaches to the searching mechanism and to the scoring function[35]. Reliability of the *in silico* methods increases when it is possible to employ different tools in order to apply a consensus for a given answer. By definition, *in silico* approaches are simulations of reality and therefore will never be completely accurate. When more than one algorithm and approach are combined, more accurate predictions are possible. If two programs that use different data and approaches are consistent, the reliability of the result is better. The errors of a given algorithm/approach must be different from the other algorithm/approach, thus being compensated as the results converge.

All simulations were carried out in triplicate. Targets (RNAp_wt, RNAp_INS1 and RNAp_INS2) and ligand (RIF) were prepared for docking simulations with the AutoDockTools (ADT)[36] interface, version 1.5.6. In all cases, the polymerase was treated as rigid and the ligand (RIF) as flexible. The grid box was centered on the C□ of the Gln403 amino acid residue of the beta subunit of RNAp. Gasteiger[37] partial charges were calculated after addition of all hydrogens. Nonpolar hydrogens of enzyme and ligand were subsequently merged.

For Vina, a protocol consisting of a cubic box of the 21 × 21 × 21 Å was established. An exhaustiveness of 8 was chosen and the other parameters kept standard. The lowest docking-energy conformation of the set of poses generated by each docking simulation was chosen for analysis.

For AutoDock, a protocol consisting of a cubic box of 80 × 80 × 80 points with a spacing of 0.35 Å between the grid points was used. Global search Lamarckian genetic algorithm (LGA)[38] and local search (LS) pseudo-Solis and Wets[39] methods were applied in the docking search. Each single docking simulation consisted of 100 independent runs. The initial population was 150, the maximum number of generations was 27,000 and the maximum number of energy evaluations was 2.5 × 106. Default values were selected for other parameters. The resulting docked conformations were clustered into families according to the RMSD. The lowest docking-energy conformation of the cluster with the lowest energy was chosen for analysis. All molecular figures were generated using Chimera software (Pettersen et al., 2004).

### 2.8. Ethics

This study was approved by the Research Ethics Committee of the Fundação Estadual de Produção e Pesquisa em Saúde (FEPPS/RS), protocol number 1.587.621 CAAE: 18269313.0.0000.5320.

## 3. RESULTS

### 3.1. Population and phenotypic drug resistance

Among the 174 RIF-resistant strains from Rio Grande do Sul, collected from 2011 to 2014, we identified 16 (9.19%) strains carrying on a 12nt insertion at codon 435 of the *rpoB* gene, all of them MDR. Amid those 16 MDR strains, one was additionally resistant to EMB and other to STR, according to DST results. Regarding the clinical and epidemiological characteristics, 14 were male, 14 previously treated for TB and seven HIV-infected. Besides that, unfavorable outcome was seen for 13 patients, the most of them due to treatment abandonment (7/16), failure to current treatment (5/16) and death in one case (Supplementary Table 1).

Nine individuals were prison inmates or spent some time in prison, and two individuals were homeless. The inmate individuals were from four different prison establishments, including the main state prisons: Cadeia Pública de Porto Alegre, Penitenciária Estadual do Jacuí and Penitenciária Modulada Estadual de Charqueadas.

### 3.2. Molecular characterization of the 12nt insertion at *rpoB* gene

Analyzing the *rpoB* gene nucleotide sequence we identified a small difference in the occurrence of the 12nt insertion. Overall, the 12nt insertion occurred at the genomic position 761111 corresponding to codon 435 of the *rpoB* gene. However, 12 of the 16 strains with the insertion presented an additional change from adenine to guanine at the position 761110, resulting in the amino acid substitution at codon 435 (GAC>GGC, Asp435Gly). To an easy comprehension, we named the 12nt insertion with additional polymorphism at genomic position 761110 of *“rpoB* INS1”, and that with only the 12nt insertion of *“rpoB* INS2” (Figure 1). None of the strains with the 12nt insertion had additional variants inside RRDR.

**Figure 1.**
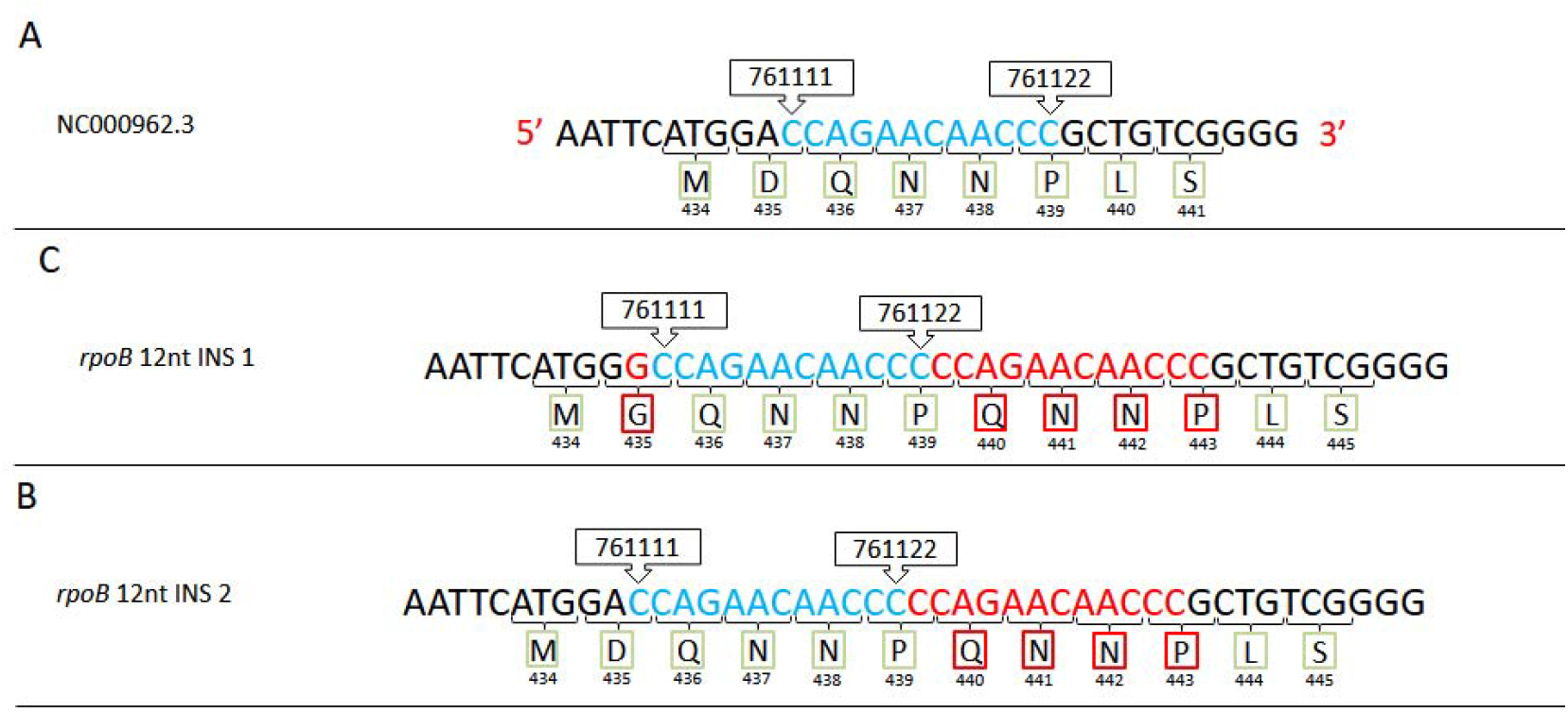
Illustration showing the occurrence of the 12nt insertion. A) wild-type sequence, B) insertion *rpoB* INS1 and C) insertion *rpoB* INS2. The numbers above the sequence refer to the nucleotide genomic position. Bellow of the sequence is annotated the amino acid corresponding to each codon and the codon number on the *rpoB* gene. Colored in blue: nucleotides from wild-type sequence; green: amino acids from wild-type sequence; and red: nucleotides and amino acids resulting from 12nt insertion.

### 3.3. Genetic diversity and phylogenetic analysis

All strains presenting the 12nt insertion were classified as belonging to *M. tuberculosis* lineage 4, sublineage 4.3.3 and to Euro-American family LAM by SNP-based typing. In the same way, all strains bear the RD115 deletion. *In silico* spoligotyping revealed that 15 strains shared the same spoligo pattern, belonging to Spoligo Internacional Type (SIT) 863, previously wrongly identified as *Mycobacterium pinnipedii*[6]. The remaining strain had absence of the 43 spacers within direct repeat (DR) locus and classified as ATYPIC (SIT 2669) on SITVIT2 database.

Besides that, phylogenetic inference including genomes from the seven *M. tuberculosis* lineages, Lineage 4 sublineages and other four SIT 863 strains with absence of the 12nt insertion, showed the closeness of strains with 12nt insertion with other SIT 863 strains (Figure 2). Another important fact revealed by WGS-based phylogenomic is that applying a five SNPs threshold, the 16 strains were grouped into a unique genomic cluster, with a mean pairwise distance of 8.94 SNPs (Figure 3).

**Figure 2.**
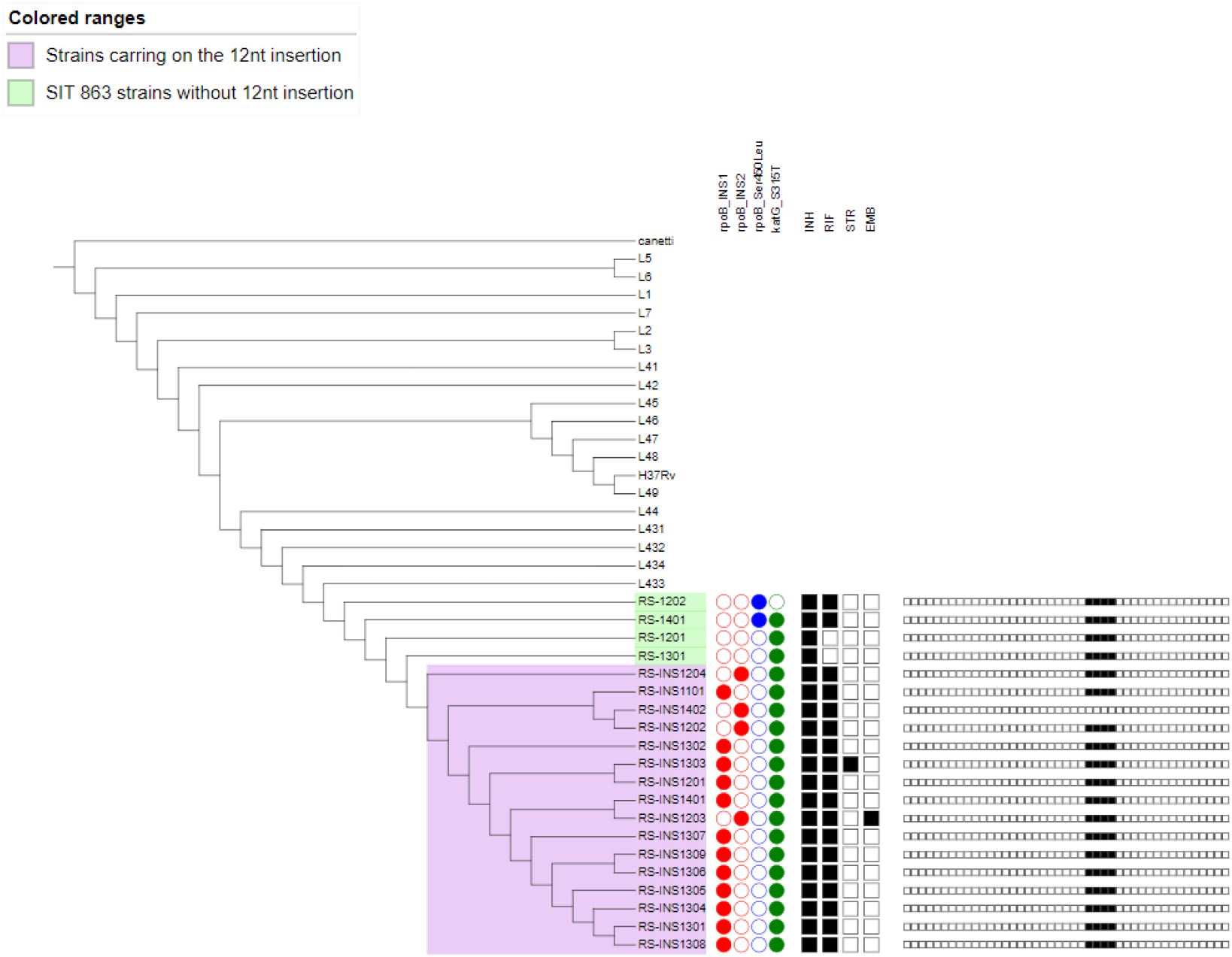
Core-genome phylogenetic tree of the 16 strains bearing the 12nt insertion, four SIT863 strains without the insertion, and representative genomes from M. tuberculosis main lineages and Lineage 4 sublineages. The tree was based on 16,566 SNPs and rooted on *Mycobacterium cannettii*. The colored ranges in purple are the strains with the 12nt insertion, and in green SIT863 strains without the insertion. The presence of mutations related to INH and RIF resistance is annotated with circles (presence: filled circle, absence: empty circle) and phenotypic drug susceptibility data for first-line drugs with the squares (resistant: filled square, susceptible: empty square). Spoligotyping profile is presented on small squares, indicating the presence (filled square) or absence (empty square) of the 43 spacer sequences.

**Figure 3.**
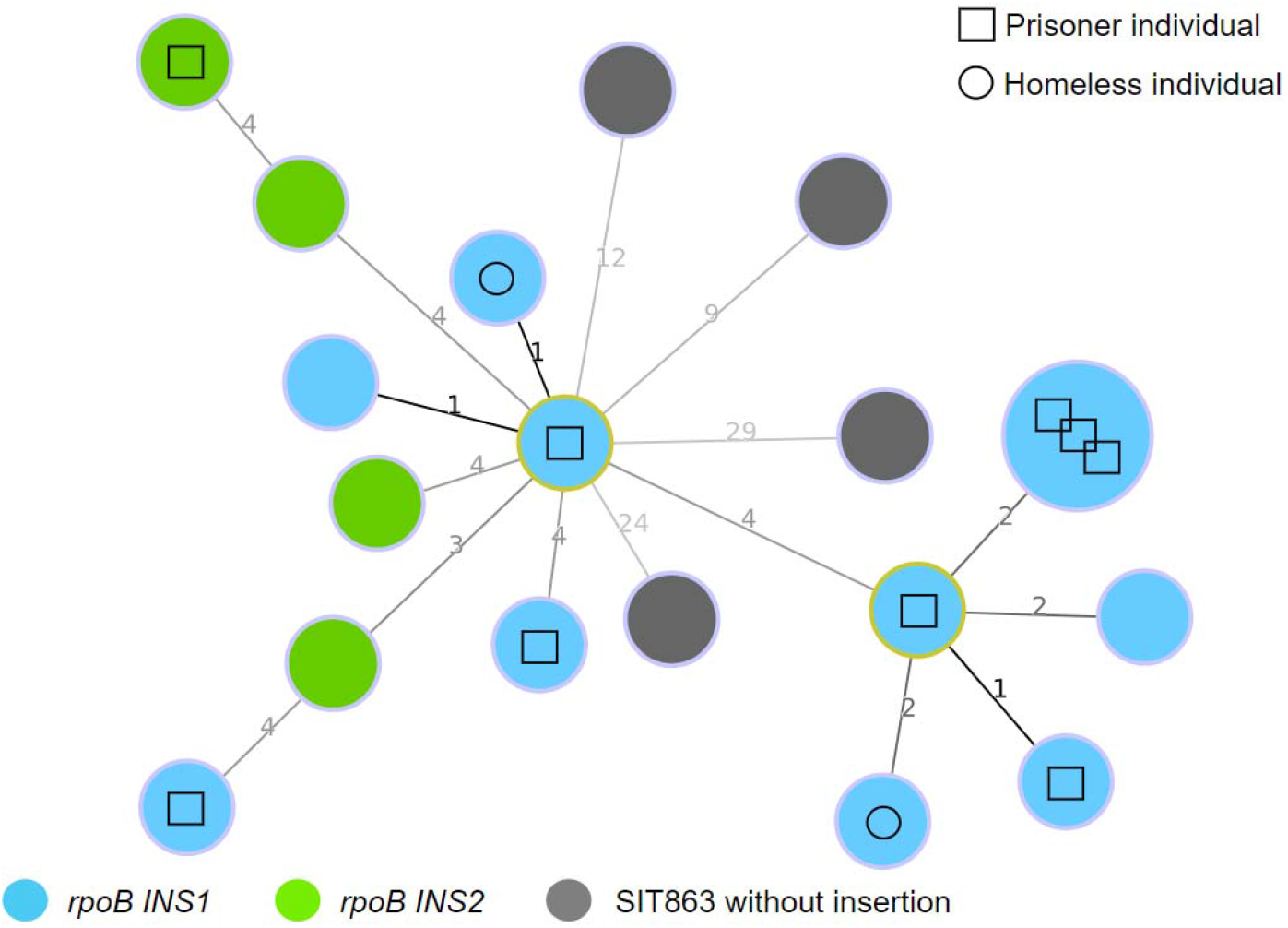
Minimum spanning tree for the 12nt insertion and SIT863 without insertion strains, based on goeBURST algorithm. The circles representing strains from prison inmate individuals are identified with a square and with a circle are flagged the strains from homeless individuals. The color of each circle indicates the insertion type (blue: *rpoB* INS1; green: *rpoB* INS2; grey: SIT863 without insertion). The number in the lines between the circles refers to the core-genome distance amid the strains. The larger circle represents three strains sharing the same core-genome.

Regarding resistance-related mutations in other genes, all the 16 strains had the Ser315Thr mutation at *katG* gene, three had variants associated to EMB resistance: one in the *embA* gene (−11C>A), one at *embB* (Met306Val) and the remaining a double mutation at *embB* gene (Gly406Asp and Met306Ile). Besides, one strain showed mutation related to ethionamide resistance (an insertion at codons 755/756 of the *ethA* gene). We also looked for putative compensatory mutations in the genes *rpoA, rpoB* and *rpoC*, the only variant identified was a synonymous substitution (Ala524Ala) at *rpoC* gene present in all the 16 strains.

### 3.4. *In silico* prediction of drug binding effect

We modeled the effect of both INS1 and INS2 on the RNAp structure in order to predict the consequence of these polymorphisms for interaction between RIF and the polymerase. At first, we performed the homology modeling followed by model evaluation using stereochemical quality analysis tools (completely showed in Supplementary Information. With the homology models validated, we used docking programs to predict RIF interaction in the modeled structures. In Table 1, it is possible to observe that the docking into the RNAP_wt binding site indicates greater affinity for the RNAP_wt-RIF interaction (average values −8.6 ± 0.0 for Vina and −9.1 ± 0.0 for AutoDock) in comparison with RNAP_INS1-RIF interaction (average values −6.9 ± 0.0 for Vina and −5.4 ± 0.3 for AutoDock) and RNAP_INS2-RIF interaction (average values −6.7 ± 0.0 for Vina and −6.1 ± 0.1 for AutoDock). Figure 4 allows evaluating the structural effect of insertions on the RIF binding site.

**Figure 4.**
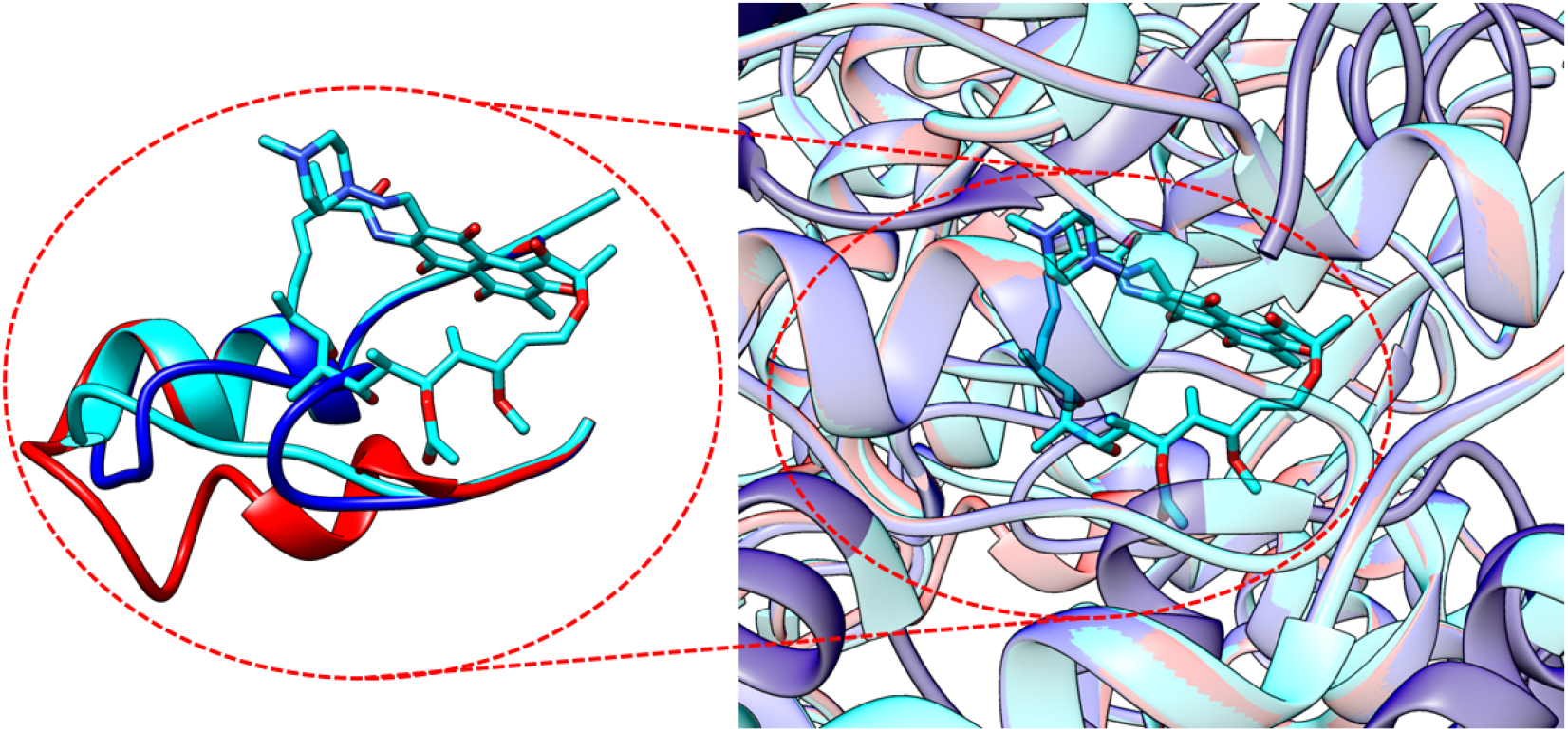
Schematic representation of the fitting of RNAP_wt (light blue), RNAP_INS1 (red) and RNAP_INS2 (navy blue) structures with emphasis on the RIF binding site. RIF is shown in its native position.

**Table 1.** Binding scores of RIF interaction with wild-type, INS1 and INS2 RNAp

| Target | Run | Binding score<br>Vina | Binding score<br>Autodock |
| --- | --- | --- | --- |
| RNAP_wt | 1 | -8.6 | -9.1 |
|  | 2 | -8.6 | -9.1 |
|  | 3 | -8.6 | -9.1 |
| RNAP_INS1 | 1 | -6.9 | -5.2 |
|  | 2 | -6.9 | -5.8 |
|  | 3 | -6.9 | -5.3 |
| RNAP_INS2 | 1 | -6.7 | -6.0 |
|  | 2 | -6.7 | -6.2 |
|  | 3 | -6.7 | -6.0 |

### 3.5. Resistance level and biological cost

All the nine strains bearing the 12nt insertion and available for additional phenotypic tests presented a MIC value ≥ to 32 mg/L. The growth index average (GI) was determined for two individual clinical strains with the 12nt insertion (GI=38.5h), for three wild type clinical strains (GI=20h) and in one susceptible control strain (H37Rv) (GI=15). The insertion strains grow 1.95 and 2.5 slower than the wild type and H37Rv, respectively, showing a fitness disadvantage.

## 4. DISCUSSION

Rio Grande do Sul is a high burden TB state in Southern Brazil with elevated rates of MDR-TB. The state has an estimated population of 11,3 million people[40] and in 2018 were accounted for 4,541 new TB cases[41] and around 82 MDR diagnosed (data from LACEN-RS). Our group has been exploring *M. tuberculosis* genetic diversity and its transmission across Rio Grande do Sul State in the last two decades. In 2012, Perizzolo *et al*.[5] first described the presence of a 12nt insertion at *rpoB* gene among multi-drug resistant strains collected between 2004 and 2006. In 2015, Dalla Costa *et al*.[6] described six strains presenting the same insertion, all of them belonging to the SIT 863 and isolated in 2006 and 2010. Latest studies in the region have shown an increased frequency of strains carrying on this insertion[7,9,42], flagging its spreading and ongoing transmission. Currently, we are performing a population-based study on drug-resistant *M. tuberculosis* strains from Rio Grande do Sul, isolated between 2011 and 2014, and identified 16 MDR strains harboring the 12nt insertion.

WGS-based typing in concert to phylogenetic analysis corroborates with previous data[6] that associated the occurrence of the 12nt insertion with strains belonging to SIT 863 that presents an important and restricted distribution in Rio Grande do Sul State[9,42]. According to Dalla Costa et al.[6], strains showing the SIT 863 pattern, and previously wrongly identified as *Mycobacterium pinnipedii*, are in fact *M. tuberculosis* Lineage 4, subfamily LAM strains. Our results are in accordance with that; all strains were classified to sublineage 4.3.3, LAM subfamily and presented RD115. These data are also confirmed by phylogenetic inference here conducted that showed the genomic closeness between 12nt insertion and SIT 863 without the insertion strains, as well as, with 4.3.3 sublineage strains. Although previous studies[5,42] related the presence of 12nt insertion strains classified as other sublineages, the genotyping techniques previously used, such as RFLP, MIRU-VNTR and spoligotyping, could produce some erroneous classification.

Aiming to identify the transmission scenario among those strains we used a five SNP genome-wide cut-off to define recent transmission, and infer that strains within that maximum genomic distance are likely epidemiologically linked[43]. The 16 strains harbouring the 12nt insertion at *rpoB* comprised a unique genomic cluster, indicative of recent and ongoing transmission of these strains in recent years. In addition, two SIT 863 strains without the 12nt insertion had a pairwise distance of 12 SNPs or less from strains bearing the 12nt insertion, corroborating with the previously cited relationships among these isolates.

The strains that presented the 12nt insertion were mostly from inmate individuals (56.25%) from different prison establishments, including the three main state prisons that are wisely overcrowded structurally limited establishments. Along to these imprisonment conditions, we observed among the individuals included in this study, the passage of the same inmate individual through different prison establishments in a short period of time, and this fact can be played an important role in the spreading of the 12nt insertion strain. Besides that, the high number of individuals that abandoned the anti-TB treatment (43.8%) and other risk factors as the homeless condition can be listed as possible in charge for the substantial ongoing transmission of this strain. However, the strains from individuals without imprisonment history within the same cluster those strains from inmate individuals point toward a transmission chain involving the inmate population and the community. The fact that the inmate population may act as a reservoir feeding the TB transmission into the community is well established[44], and requires attention in the public health measures that aim to control the TB transmission in different settings.

In order to obtain a better understanding regarding the 12nt insertion occurrence, we accessed *rpoB* gene nucleotide sequence and observed that 12nt insertion occurs with a discrete variance amidst strains. Thus, we obtained RNAp structures carrying on both *rpoB* INS1 and *rpoB* INS2 structures by homology modeling, and docked those structures, as well as the RNAP wt structure with the RIF molecule. The decreased affinity of RIF for the RNAp_INS1 and RNAp_INS2 mutant forms may explain the resistance of the strains that carry the mutations. It is also possible to observe that the insertions promote changes in the binding site, which is probably linked to the lower affinity of RIF interaction with the mutant forms.

We explored the phenotypic impact of the 12nt insertion concerning resistance level to RIF and to growth rate of the *M. tuberculosis* strains. Using the resistance level classification defined in the classical study of Huitric *et al*[45], the nine clinical isolates tested showed a high level of resistance (MIC> 32 μg/mL) to RIF. As stated *in silico* analysis, the insertion of the 12 nucleotides probably causes a low affinity of RIF to its target in the β-subunit of RNA polymerase coding by the *rpoB* gene and reflects in the resistance level. In relation to the fitness, we observed that the two isolated tested presented a slower growth index when compared to wild type clinical isolates and the standard H37Rv. Considering that *rpoB* is a subunit of the RNA polymerase enzyme responsible for transcription from DNA to RNA, which is an essential process in the duplication of bacteria, a further change in its structure may have influenced in this process and consequent growth time. Previous studies showed biological cost associated to rpoB mutations[46,47], however it could be mitigated with compensatory mutations[48]. It is important to point out that spite of the 12nt insertion may cause a biological cost, is difficult to extrapolate this phenotypic characteristic to an epidemiologic setting, since it is an interplay of environmental, pathogen and host factors.

## 5. CONCLUSION

Here we described the key molecular and phenotypic aspects from a *M. tuberculosis* strain emerging in Southern Brazil in recent years that carries an uncommon 12nt insertion at *rpoB* gene leading high-level RIF-resistance. The phylogenetic approach demonstrated a potential transmission from the prison inmate population to community of the 12nt insertion strain, demonstrating an urgent need for enhanced surveillance on TB transmission amid the inmate population and the anti-TB treatment abandonment, especially for MDR, in this region. Due to the high level of resistance to RIF - the most important drug in anti-TB treatment - and its established ongoing transmission, it is essential to monitor this strain spreading among the population. Since this is a retrospective study and included clinical samples collected until 2014, it is necessary to take up a more fast and, rather, real-time surveillance for this described strain in order to break up its transmission. For this propose, it would be necessary a rapid PCR-based test, for a fast, easy and low-cost identification of this strain.

## Acknowledgments

We are grateful to TGen, C-Path and ReSeqTB for supporting whole genome sequencing and to Brazilian Network of Tuberculosis Research (REDE-TB) for enabling this partnership. The authors would like to thank Centro de Desenvolvimento Científico e Tecnológico (CDCT)/CEVS/SES/RS for the support and infrastructure.

## Data availability

*Mycobacterium tuberculosis* genome data were deposited in the NCBI under BioProject IDs: PRJNA 535343 and PRJNA639713, see supplementary spreadsheet.

---

**Supplementary Table 1.** Clinical and epidemiological characteristics from 16 individuals infected by *rpoB* 12nt insertion *M. tuberculosis* strain

| Sample | Patient type | Outcome | Gender | Age (years) | HIV | Prison inmate | Homeless |
| --- | --- | --- | --- | --- | --- | --- | --- |
| RS-INS1101 | After abandoning previous treatment | Lost to follow-up | M | 33 | Neg |  | yes |
| RS-INS1201 | After abandoning previous treatment | Lost to follow-up | M | 33 | Neg | yes |  |
| RS-INS1202 | New case | Missing data | M | 44 | Pos | Missing data | Missing data |
| RS-INS1203 | New case | Cured | F | 55 | Pos |  |  |
| RS-INS1204 | New case | Treatment failure | M | 55 | Positive |  |  |
| RS-INS1301 | After abandoning previous treatment | Lost to follow-up | M | 35 | Neg | yes |  |
| RS-INS1302 | Relapse | Treatment failure | M | 39 | Pos | yes |  |
| RS-INS1303 | After abandoning previous treatment | Lost to follow-up | M | 25 | Pos | yes |  |
| RS-INS1304 | Treatment after previous treatment failure | Treatment failure | M | 43 | Neg | yes |  |
| RS-INS1305 | After abandoning previous treatment | Died | M | 33 | Pos |  |  |
| RS-INS1306 | Relapse | Cured | M | 39 | Neg | yes |  |
| RS-INS1307 | Treatment after previous treatment failure | Lost to follow-up | M | 41 | Pos |  | yes |
| RS-INS1308 | After abandoning previous treatment | Lost to follow-up | M | 25 | Neg | yes |  |
| RS-INS1401 | Treatment after previous treatment failure | Treatment failure | M | 34 | Neg | yes |  |
| RS-INS1402 | Treatment after previous treatment failure | Treatment failure | F | 43 | Neg |  |  |
| RS-INS1309 | After abandoning previous treatment | Lost to follow-up | M | 24 | Neg | yes |  |

